# Effect of gedunin on glioblastoma proliferation and invasiveness: *in vitro* and *in vivo* approaches

**DOI:** 10.1101/2023.06.02.543101

**Authors:** Thadeu Estevam Moreira Maramaldo Costa, Tatiana Almeida Pádua, Jonathas Xavier Pereira, Vinicius Gonçalves Rodrigues, Leonardo Noboru Seito, Erika Marques da Cunha, Claudio C. Filgueiras, Yael Abreu-Villaça, Alex C. Manhães, Clarissa Menezes Maya-Monteiro, Maria das Graças Henriques, Thomas Eichenberg Krahe, Carmen Penido

## Abstract

**Background:** Glioblastoma is the most invasive and malignant brain tumor, for which no effective treatment is currently available. Heat shock protein 90 (HSP90) is a potential target for the treatment of different types of cancer. Herein, we investigated the effect of gedunin, an HSP90 inhibitor, in murine GL261 glioblastoma *in vitro* and *in vivo* models.

**Methods:** Gedunin effects were assessed by *in vitro* cell viability assay (MTT), apoptosis (PI/Annexin staining), proliferation (CFSE), invasion (transmigration, scratch assay and zymography), protein detection (western blot) and *in vivo* orthotopic glioblastoma model (brain computed tomography and histology).

**Results:** Gedunin treatment decreased GL261 cell proliferation and triggered apoptosis in a concentration-dependent manner. Gedunin also reduced GL261 cell migration and metalloproteinase-2 activity, suggesting that it impairs glioblastoma cell invasion. Despite the reduction of total protein content in gedunin-treated cells, the phosphorylation of STAT3 and ERK1/2 pathways was enhanced within 24 h. *In situ* treatment with gedunin did not significantly reduce tumor volume. Still, it reduced the tumor central area and the presence of vascular structures in xenograft glioblastoma in C57BL/6 mice, a clinical feature associated with morbidity and mortality.

**Conclusion:** Gedunin inhibits murine glioblastoma cell growth and proliferation by inducing apoptosis.

**Graphical Abstract:** 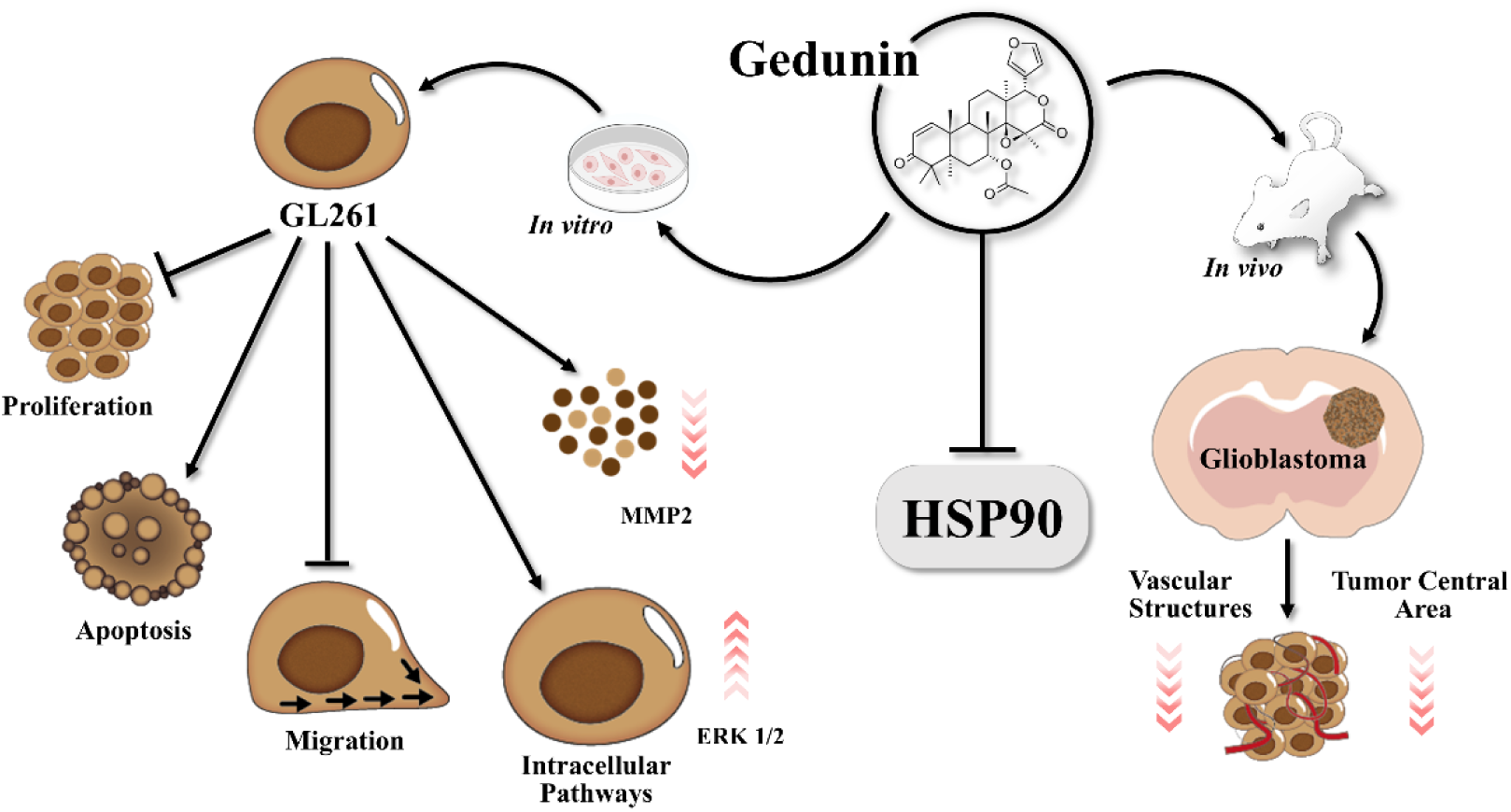

## 1. Introduction

Glioblastoma is a highly invasive and malignant tumor defined by the World Health Organization as grade IV, based on histological, immunohistochemical, and molecular features [1]. Patients bearing glioblastoma have a poor prognosis, with estimated median survival rates of eight months and a maximum five-year survival rate of only 5% [2,3].

The current standard protocol for glioblastoma treatment relies on total tumor excision followed by radiotherapy with concurrent and adjuvant chemotherapy with temozolomide [4]. The most common adverse events related to temozolomide treatment are myelosuppression and hepatotoxicity [5–7]. Another limiting factor of glioblastoma treatment is that most orally and intravenously administered alkylating agents induce resistance and have poor blood-brain barrier permeability [8,9]. An alternative strategy is the usage of intracranial wafer implants containing the alkylating agent carmustine after tumor removal surgery, which, in combination with radiochemotherapy, has been shown to significantly improve patient survival [10,11]. However, carmustine wafers induce severe side effects, such as cerebral edema and hematological toxicity [5,12,13]. Glioblastoma is still considered incurable [14]; however, treatments targeting multiple signaling pathways are a promising strategy [15].

Modulation of the chaperone heat shock protein 90 (HSP90) is considered a potential antitumor therapy, including gliomas [16–18]. HSP90 activity is implicated in several hallmarks of cancer, such as proliferation, invasion, metastasis, angiogenesis, and resistance to apoptosis [19]. Increased expression of HSP90 correlates with tumor malignancy and poor prognosis in different types of cancer, including breast, colon, and melanoma [20–23]. Indeed, HSP90 chaperones several proteins involved in oncogenic processes, including the human epidermal growth factor receptor 2 (HER2/neu; ErbB2), tyrosine-protein kinase Src (Src), protein kinase B (AKT/PKB), breakpoint cluster region/Abelson murine leukemia virus (BCR/ABL) and tumor protein 53 (p53) [24–26]. In line with this, encouraging pre-clinical data have been obtained with HSP90 inhibitors in different types of tumor cells and animal models, including lymphoma, melanoma, breast, and lung cancer [27–31]. Currently, in the National Institutes of Health (NIH) Clinical Trials database at ClinicalTrials.gov, there are more than a hundred registered trials (11 currently active) evaluating HSP90 inhibitors (alone or in combination with other drugs) for the treatment of different types of cancer [32]. Worthy of note, off-target toxicity, including ocular toxicity and hepatotoxicity, is a limiting factor for the use of these substances [33,34].

Over the past two decades, our group and others have been studying the antitumoral and anti-inflammatory effects of gedunin, a limonoid produced by vegetal species from the Meliaceae family [35] or by semi-synthetic routes [36]. Gedunin negatively modulates HSP90 activity by binding to the HSP90 co-chaperones p23 and cdc37 and, thus, impairing the formation of the active HSP90 complex [37,38]. This limonoid has potent antitumoral effects demonstrated against an array of tumor types *in vivo* and *in vitro* [37,39–42]. Gedunin anti-tumoral effect on glioblastoma has been shown *in vitro* human cells lines U87 and U251 [43,44]. However, this effect has been observed in only a few studies using cell lines. Therefore, further efforts are needed to reproduce previous findings and to further investigate the anti-glioblastoma effect of gedunin in an *in vivo* model. In the present study, we used the murine GL261 cell line in *in vitro* assays to translate the data obtained *in vitro* to a syngeneic *in vivo* model, using the well-established orthotopic *in vivo* glioblastoma model.

## 2. Materials and Methods

### 2.1 Reagents and antibodies

Gedunin (Gaia Chemicals, USA) and the specific HSP90 inhibitor 17-AAG (17-N-allylamino-17-demethoxygeldanamycin, Tanespimycin, Sigma-Aldrich, USA) were dissolved in DMSO (dimethyl sulfoxide, Sigma-Aldrich) and frozen at −86 °C until use. MTT (3-(4,5-dimethylthiazol-2-yl)-2,5-diphenyl tetrazolium bromide), Dulbeccós modified Eaglés medium (DMEM), phosphate buffer saline (PBS) tablet, and trypsin-EDTA (ethylenediamine tetraacetic acid) solution were purchased from Sigma Aldrich. Fetal bovine serum (FBS) was purchased from Cultilab (Brazil). Propidium iodide (PI) solution was obtained from Biolegend (USA). Vybrant^™^ CFDA-SE (carboxyfluorescein diacetate succinimidyl ester) Cell Tracer Kit and gentamicin were obtained from Thermo Fischer Scientific (USA). Annexin V-FITC (fluorescein isothiocyanate) conjugated was purchased from BD Pharmingen (USA). Antibodies against rabbit p44/42 MAPK (mitogen-activated protein kinase, Thr202/Tyr204, D13.14.4E), ERK1/2 (extracellular signal-regulated kinase 1/2, rabbit) and GAPDH (glyceraldehyde-3-phosphate dehydrogenase mouse, D4C6R) were purchased from Cell Signaling (USA). Anti-rabbit-IgG-HRP (horseradish peroxidase) secondary antibody was purchased from Santa Cruz Biotechnologies (USA). DNase I recombinant RNase free was purchased from Roche Life Sciences (Germany).

### 2.2 GL261 cell culture and treatment

Murine glioblastoma cell line (GL261, Cell Bank from Federal University of Rio de Janeiro - UFRJ, Brazil) was cultured in DMEM (Sigma Aldrich) supplemented with 10% FBS (Cultilab) and 50 μg/mL gentamicin (Thermo Fisher Scientific) in humidified atmosphere (5 % CO_2_ at 37 °C) until confluence. Cells were digested with 0.25 % trypsin at the 5^th^ passage and washed with 1 × PBS. Cell viability was determined by trypan blue (Sigma Aldrich) staining under a light microscope, before further assays. GL261 cells were incubated with 6.25-100 µM of gedunin for 24-72 h at 37 °C, 5 % CO_2_.

### 2.3 Animals

Adult male C57BL/6 mice (20-25 g) provided by Oswaldo Cruz Foundation (Fiocruz) breeding unit (Institute for Biomodels Science and Technology, Rio de Janeiro, Brazil) were kept in standard autoclavable polypropylene boxes, with free access to food and fresh water in environmentally controlled room temperature (ranging from 22 to 24 °C) on a light/dark cycle at Farmanguinhos (Fiocruz, Rio de Janeiro, Brazil) and at the Laboratory of Neurophysiology (State University of Rio de Janeiro, Rio de Janeiro, Brazil) experimental animal facilities. Animal procedures were performed in accordance with the ethical guidelines of the Brazilian National Council for the Control of Animal Experimentation (CONCEA) and reported following the ARRIVE guidelines [45]. Experimental procedures were approved by the Ethics Commission on Animal Use of Oswaldo Cruz Foundation (license #LW-35/16) and the Ethics Commission on Animal Use of the State University of Rio de Janeiro (license #025/2017).

### 2.4 Primary mixed glial cell culture

Under sterile conditions, mouse brain cortex tissue was rapidly dissected from euthanized (150 mg/kg sodium thiopental, i.p.) young adult (4-week-old) C57BL/6 male mice and transferred to ice-cold DMEM/F12 media (Sigma Aldrich) in 6 well plates. After removal of the meninges, brain tissue was cut into small pieces, and washed three times with 1x sterile PBS (Sigma Aldrich). Trypsin-DNAse solution (0.25 % - 250 µg/mL) was added to the wells and the plate was incubated for 1 h in an orbital shaker (Excella E24, New Brunswick Scientific, USA) at 37 °C and 150 rpm. Tissue samples were vigorously triturated and dissociated with a 1 mL pipette tip. Cells were washed with FBS-DNAse solution (10 % - 250 µg/mL), added to a 20 % percoll gradient media, and centrifuged at 800 × *g* for 30 min at 25 °C. Recovered cells were washed (400 × *g*, 10 min at 4 °C), distributed in T-25 bottle flasks, and incubated at 37 °C, 5 % CO_2_. DMEM/F12 media was changed every 3-4 days until use. Data were collected from two independent experiments (n=5 mice per group; license #LW-35/16).

### 2.5 Proliferation assay

GL261 cells (4 × 10^7^/mL) were incubated with 5 μM CFSE solution (Invitrogen Molecular Probes, USA) diluted in PBS for 15 min at 37 °C, 5 % CO_2_. Cells were washed with 10 % PBS/FBS solution, seeded in 24 well plates (2 × 10^5^ cells per well), incubated overnight, and treated with gedunin (6.25-100 μM) for 72 h. Cell proliferation was evaluated by analyzing CFSE fluorescence intensity (FL1) upon cell divisions, by flow cytometry (FACScalibur, Becton Dickinson, USA). Results were analyzed using FlowJo X Software (Tree Star, Becton Dickinson).

### 2.6 MTT assay

GL261 or primary mixed glial cells were seeded in a 96 flat bottom well plate (2 × 10^4^ cells/well) and cultured in DMEM for 24 h (CO_2_ 5 % at 37 °C) for adhesion. Adherent cells were cultured with different concentrations of gedunin (6.25-100 μM), 17-AAG (1 μM), or temozolomide (100 μM) for 20 or 68 h, when MTT solution (5 mg/mL; 22.5 μL/well) was added to the culture and incubated for 4 h. The analyzes were performed at 24 and 72 h time points. Plates were centrifuged for 3 min (720 × *g* at 4 °C), the supernatant was discharged and DMSO (150 μL/well) was added for formazan crystals solubilization. The absorbance was read in a microplate reader (SpectraMax M5, Molecular Devices, USA) at 540 nm.

### 2.7 Determination of apoptosis

Apoptosis was evaluated using annexin V-FITC and propidium iodide (PI) staining. GL261 cells were seeded in 24 well cell culture plates (2 × 10^5^ cells/well) and left to adhere overnight. GL261 cells were then incubated with gedunin (6.25 - 100 μM) for 24 or 72 h, followed by trypsinization, resuspension in annexin V Binding Buffer (HEPES 10 mM pH 7.4, NaCl 140 mM, CaCl_2_ 2.5 mM) and labeling with annexin V-FITC and PI for 15 min at room temperature. Cells incubated with medium were used as the negative control, whereas 17-AAG-incubated cells were used as the positive control. Staining was analyzed by cytometry using FACScalibur (Becton Dickinson) and FlowJo Software (Tree Star, Becton Dickinson). At least 10^4^ cells were acquired per sample. Data are presented in dot plots in a log scale of increasing FL-1 (annexin V-FITC) and FL-3 (PI) intensity.

### 2.8 Scratch assay

GL261 cell monolayers, seeded into 6 well plates, were scratched at the center of the well, and then the migration path was assessed at the leading edge of the scratch by phase-contrast optical microscopy (100 ×, Olympus IX70, Japan) at indicated time points. Analyses were performed using Image J software (National Institute of Health, USA) **[46]**. The rate of migration was determined by quantifying the total distance that cells moved from the edge of the scratch toward the center of the scratch, as previously described **[47]**.

### 2.9 Chemotaxis assay

GL261 cell migration was assayed in a 48-well micro-chemotaxis Boyden chamber (Neuro Probe Inc., USA). The bottom wells of the chamber were filled with DMEM 10% FBS (29 µL) or serum-free DMEM media as the negative control. Polycarbonate membrane filters with 8 µm pore size (Neuro Probe) were placed between the lower and upper chambers. GL261 cells were pre-treated with different concentrations of gedunin (6.25-25 μM) for 1 h in serum-free DMEM and then loaded into the upper wells of Boyden chamber (2 × 10^4^ cells/50 μL/well). The chamber was incubated in a humidified atmosphere with 5 % CO_2_ at 37 °C for 24 h. The membranes were removed and stained using a Diff-quick (Laborclin, Brazil). The bottom side of the membranes was visualized by light microscopy for counting GL261 migrating cells (1000 ×, Olympus IX70). GL261 chemotaxis was calculated and expressed as the mean number of migrated cells in ten random high-power fields (HPFs) per well (in quadruplicate).

### 2.10 Zymography

GL261 cells were plated (1 × 10^6^/well) in a 6-well plate in DMEM. After 24 h, supernatant was removed, and cells were washed 3 times with pure DMEM to eliminate any FBS trace. Then, cells received gedunin (6.25-50 μM) or 17-AAG (1 μM) treatment in serum-free DMEM. After 48 h, the supernatant was collected and stored at −70 °C until lyophilization. Samples were dehydrated in a Gamma 2-16 LSCplus (Christ, Germany) ice condenser and, resuspended with 200 μL of zymography extraction buffer (Tris-HCl 50 mM pH 7.5, NaCl 0.2 M, Triton X-100 0.1 %, CaCl_2_ 10 mM, protease inhibitor cocktail). Lowry method was used for protein quantitation. Enzymatic activity of matrix metalloproteinase (MMP)-2 was determined on a 10 % SDS-PAGE containing 0.1 % gelatin from porcine skin (Sigma Aldrich) [48].

### 2.11 Western blot

GL261 cells (1 × 10^5^ cells) treated with gedunin (6.25-100 μM) or 17-AAG (1 µM) for 24 h were lysed in a buffer containing protease inhibitors. Protein extract (20 µg) was resolved by SDS-PAGE, proteins were transferred to polyvinylidene difluoride (PVDF) membranes (GE Healthcare, USA), and then probed with primary antibodies. Antibodies were diluted in Tris-buffered saline containing 0.5 % Tween 20 and 5 % non-fat milk or bovine serum albumin (BSA, Sigma - Aldrich, USA), and incubated overnight at 4 °C. The membranes were incubated with horseradish peroxidase (HRP)-conjugated secondary IgG (Santa Cruz Biotechnology) for 1 h at room temperature. The blots were developed in a Hyperfilm with the use of a chemiluminescent substrate (ECL; Amersham Biosciences, USA). The band intensity was quantified using ImageJ (NIH, USA), and expressed in arbitrary units (a.u.). The raw intensity values of total proteins were normalized using the values of the constitutive protein (GAPDH, used as loading control) and of the control group (untreated cells) to allow comparison between different experiments. The phospho-protein values were normalized by the specific total protein content. Western blot membranes were stripped using harsh stripping buffer (SDS 10% Tris HCl, pH 6.8, 0.5 M; 0.8 % ß-mercaptoethanol), at 50 °C for up to 45 min under agitation in an orbital shaker (Excella E24, New Brunswick Scientific, USA). Stripping was performed between the phospho and total proteins antibody incubations. Analyses were performed using Image J software (NIH, USA)**[46]**.

### 2.12 Orthotopic glioblastoma model and osmotic mini-pump implantation

Glioblastoma was induced in adult male C57BL/6 by stereotaxic injections of GL261 cells at the Laboratory of Neurophysiology experimental animal facility (UERJ, Rio de Janeiro, Brazil). Mice were intraperitoneally anesthetized with 100 mg/kg ketamine (Syntec, Brazil) and 10 mg/kg xylazine (Sedalex™ Venco, Brazil). Under sterile conditions, a midline incision was made in the scalp, the skull was exposed to identify bregma and lambda, and a small burr hole was drilled over the right hemisphere. GL261 cells (5 × 10^4^/μL; final volume of 2 μL) were stereotaxically injected into the caudate putamen with a 10 μL Hamilton syringe at the rate of 0.5 μL/min. The needle was slowly removed 4 min after the injection was finished. The following coordinates (in mm) were used for stereotaxic injections (anterior-posterior; medial-lateral; dorsal-ventral): 0.1 AP; 2.32 ML; 2.35 DV. Coordinates were based on the Cha et al., 2003 [49] descriptions and confirmed on Paxinos and Franklin mouse brain atlas [50]. Fourteen days after GL261 injections, animals were implanted with Alzet osmotic minipumps (model 2004, 0.25 μL/h; Durect Corporation, USA) filled with gedunin (250 μM, n=8) or DMSO (2.5%, n=8). Cannulas (1-2 mm long) were placed into the site of the tumor following similar surgical procedures described above. Treatment lasted between 14 and 17 days. Mice were monitored every day for clinical signs of pain and distress, such as ruffled fur, hunched back, trembling, convulsions, paralysis, and permanent recumbency [51]. If signs of pain and distress were observed, the animal was excluded from the study and euthanized. Data were collected from two independent experiments (n=8 randomized mice per group). Mice were euthanized 28 days after GL261 injection with sodium thiopental (150 mg/kg, i.p.).

### 2.13 Mouse Brain Computed Tomography Images

After 14-day treatment, anesthetized mice (100 mg/kg ketamine/10 mg/kg xylazine, i.p.) were intravenously (i.v.) injected with 300 mg/mL of Omnipaque iohexol contrast (GE Healthcare). Brains were scanned 3-6 min after contrast administration in a GE Highspeed CT/E image system (Pedro Ernesto University Hospital, UERJ, Rio de Janeiro, Brazil), according to the following protocol: 120 kV tube voltage, 55 mA, 1 mm slice thickness. Images were reconstructed in a 256 x 256-pixel matrix using InVesalius 3.1.1 software (developed by Centro de Tecnologia da Informação Renato Archer, CNPq, and Ministry of Health, Brazil). Tumor volumetry was determined by selecting contrast-enhanced regions of interest in coronal sections.

### 2.14 Histology of Brain Tissue

Euthanized mice (150 mg/kg sodium thiopental, i.p.) were perfused with 4 % formaldehyde. Mouse brains were collected and transferred into a 4 % formaldehyde solution for 2 days for tissue fixation. Brains were then sectioned and embedded in paraffin. Coronal sections of brain tissue (5 μm) were made using a Shandon Finesse 325 microtome (Thermo Fisher Scientific). Brain slices were stained with hematoxylin (Merck, USA) and eosin (Sigma Aldrich, USA). Tumor central area was measured using Axioplan^®^ software [52]. Lesion size, liquefactive necrosis area, vascular structures, intratumoral hemorrhage, and mitosis were blindly analyzed in the central slice tumor area. Lesion size area was defined by the percentage of tumor area relative to the size of the total brain area in the slice. The liquefactive necrosis area was determined by the percentage of necrosis area relative to lesion size. Vascular structures with their full composition were counted in both intratumor and peritumoral regions. Intratumor hemorrhage was described in a semi-quantitative form regarding the intensity at the slice, with the following definitions: discrete (+), moderate (++), and intense (+++). Mitosis was characterized by estimating the average amount of cells per analyzed field, and thus: >50% (+), 50-75% (++), and >75% (+++). Multipoint brain slice images were composed in NIS-Elements AR (Nikon Corporation). Each brain slice was analyzed by an observer blind to the experimental groups.

### 2.15 Statistical Analysis

All analyses were performed using GraphPad Prism 8 (GraphPad Software, USA). Western blot analyses were performed using the non-parametric Kruskal-Wallis test followed by Dunnett’s multiple comparisons. Histologic analyses were performed by one-way analysis of variance (ANOVA) and Tukey’s post-test for multiple comparisons. All other comparisons were carried out using one-way ANOVA followed by the Student-Newman-Keuls test. Data are shown as the mean ± SEM (unless otherwise mentioned). Significance was assumed either at the level of p ≤ 0.05, p ≤ 0.01, or p ≤ 0.001, as specified.

## 3. Results

### 3.1 Gedunin impairs GL261 cell growth and viability in vitro

To verify the effect of gedunin on GL261 cell growth, a 72 h CFSE proliferation assay was performed, in which staining decreases as cells proliferate. Incubation of CFDA-SE-stained GL261 cells with gedunin decreased cell proliferation in a concentration-dependent manner (**Fig. 1A**). Gedunin (25-100 µM) significantly inhibited GL261 cell growth, similar to what was observed with the selective HSP90 inhibitor 17-AAG (reference control). **Figure 1B** shows representative histograms of CFSE fluorescence intensity (FL-1) of GL261 cells depicted in **Figure 1A**.

**Figure 1.**
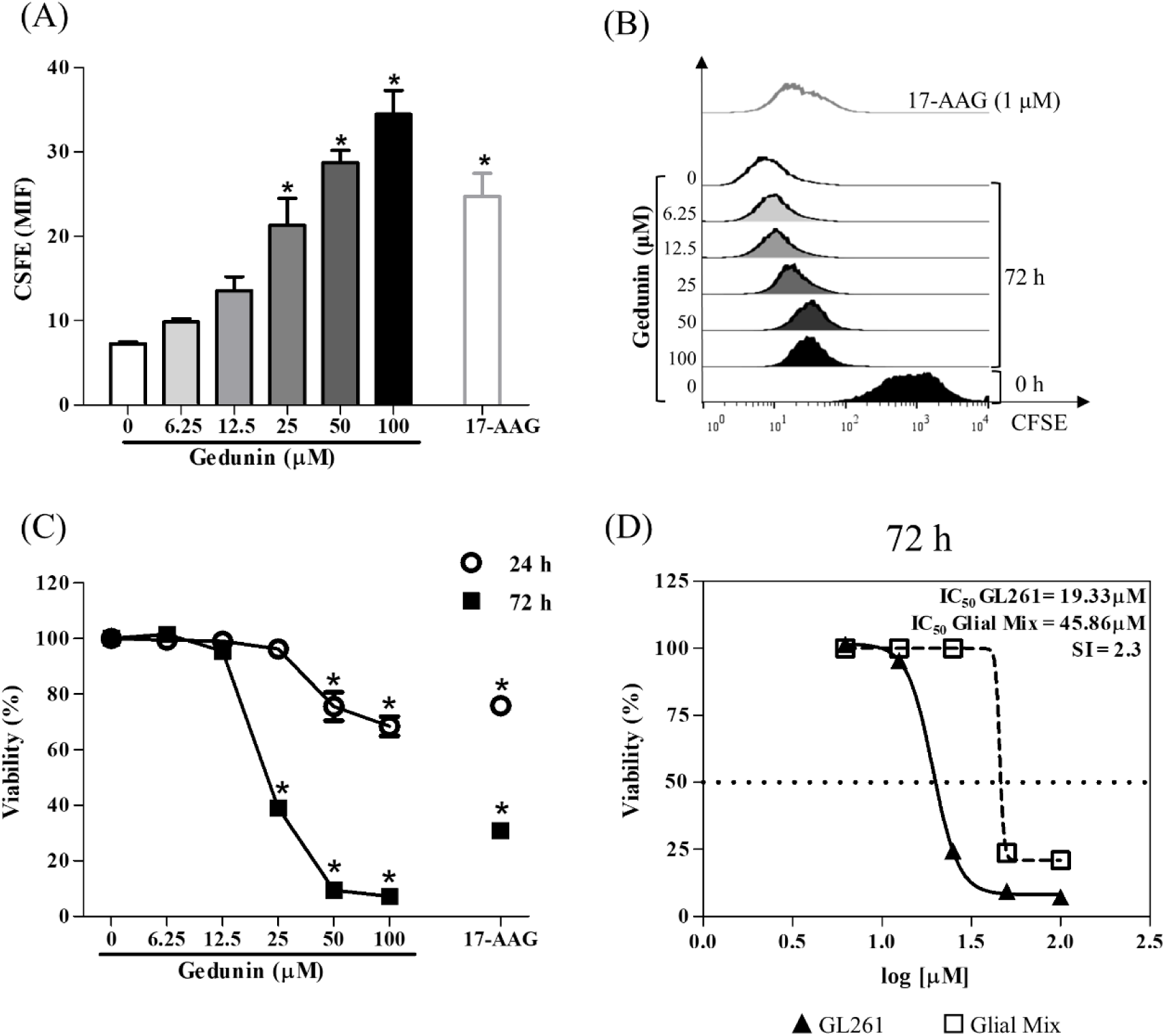
Gedunin impairs GL261 cell proliferation. (A) Effect of gedunin (6.25-100 μM) and 17-AAG (1 μM) on GL261 cell proliferation evaluated by CFSE staining and analyzed by FACScalibur (BD Biosciences) 72 h after incubation. Results are expressed as the mean ± SEM of median fluorescence intensity (MFI) from triplicate wells from three independent experiments. (B) Representative histogram shows CFSE fluorescence intensity (FL-1) from one experiment. (C) Effect of gedunin (6.25-100 μM) and 17-AAG (1 μM) on GL261 cell viability evaluated by MTT reduction assay (SpectraMax M5, Molecular Devices) 24 and 72 h after incubation. (D) IC50 values of gedunin treatment on GL261 and glial mixed cells 72 h after incubation. SI: selectivity index.

The effect of gedunin on cell viability was evaluated by the MTT assay. The incubation of GL261 cells with gedunin decreased mitochondrial function in a concentration-dependent manner within 24 and 72 h (**Fig. 1C**). MTT assay was also performed on primary mixed glial cells to evaluate gedunin toxicity to non-tumoral cells. Gedunin effect on primary mixed glial cells was less toxic, and higher concentrations were required to induce cell death when compared to GL261 cells (**Fig. 1C**). The half-maximal inhibitory concentration [IC50] of gedunin on GL261 cells was 2.3 times lower than the IC_50_ observed on primary glial cells (**Fig. 1D**). Temozolomide, the first-line treatment for glioblastoma, also reduced GL261 viability within 72 h in a concentration-dependent manner (**Suppl. Fig. 1**). Considering the antiproliferative and cytotoxic effects of gedunin on GL261 cells, we further examined whether this limonoid induced apoptosis.

### 3.2 Gedunin induces GL261 cell apoptosis in vitro

Apoptosis is regulated by a series of signaling events that include disruption of the mitochondrial membrane and phosphatidylserine externalization [53,54]. Membrane phosphatidylserine staining by annexin V-FITC revealed that gedunin induced apoptosis of GL261 cells in a concentration- and time-dependent manner (Fig. 2A). Apoptosis was detected 24 h after treatment, and surpassed 90 % within 72 h, at the highest concentrations. Representative dot plots of PI^+^ and annexin V^+^ cells are shown in Figure 2B. Apoptotic cells are characterized by reduced forward scatter and increased side scatter [55–57]. Indeed, GL261 cells incubated with gedunin diminished FSC-H and increased SSC-H intensities, when compared to non-treated cells (Fig. 2C), supporting results shown in Figures 2 A and B. Representative dot-plots of the effect of 17-AAG (1 µM) on GL261 apoptosis (24 h and 72 h) is available on **Supplemental** Figure 2.

**Figure 2.**
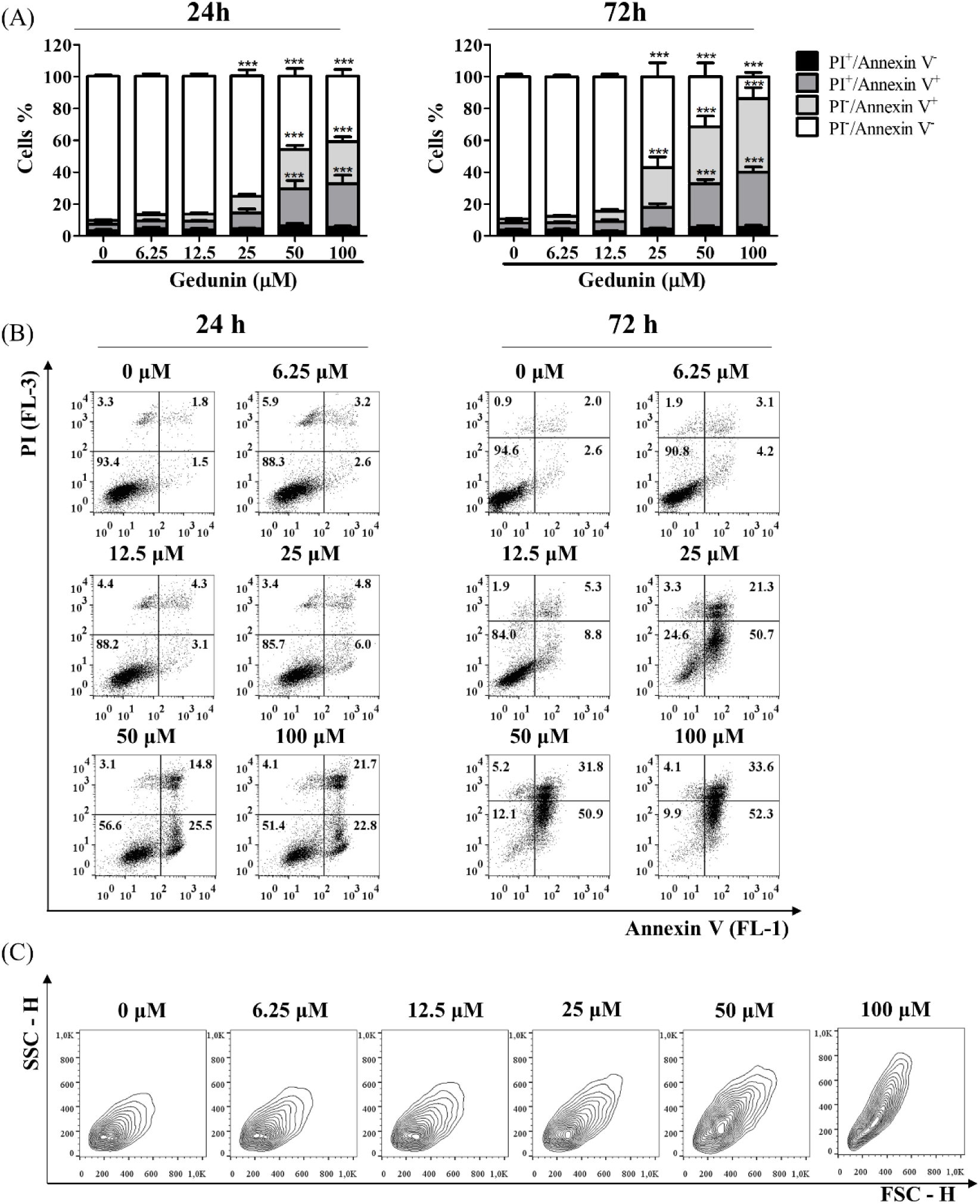
Gedunin induces time- and concentration-dependent GL261 apoptosis. (A) Effect of gedunin (6.25-100 μM) on GL261 cell (2 × 10^4^/well) apoptosis evaluated by propidium iodide and annexin V-FITC staining MTT (FacsCalibur – BD Biosciences) 24 h and 72 h after incubation. Results are expressed as the mean ± SEM of viability percentage from triplicates wells from three independent experiments (24 well plates). * indicates statistically significant differences (p ≤ 0.05) between treated and untreated cells. (B) Representative dot plots of flow cytometry analysis of gedunin-treated GL261 cells labeled with annexin V-FITC and PI after 24 and 72 h incubation. (C) Representative contour plots of SSC-H and FSC-H of gedunin-treated (6.25 - 100 μM) GL261 cells after 72 h.

### 3.3 Gedunin impairs GL261 cell migration and invasiveness in vitro

Tumor cell migration is an earlier step in cancer dissemination and invasion [58,59]. Scratch assay shows that gedunin (12.5 and 25 µM) reduced GL261 cell migration when compared to the untreated group within 24 h, 48 h, and 72 h (Fig. 3A, B). Confirming results obtained in the scratch assay, reduction of chemotaxis was also observed in the Boyden chamber migration assay, in which gedunin treatment (25 µM) impaired cell migration within 48 h (Fig. 3C).

**Figure 3.**
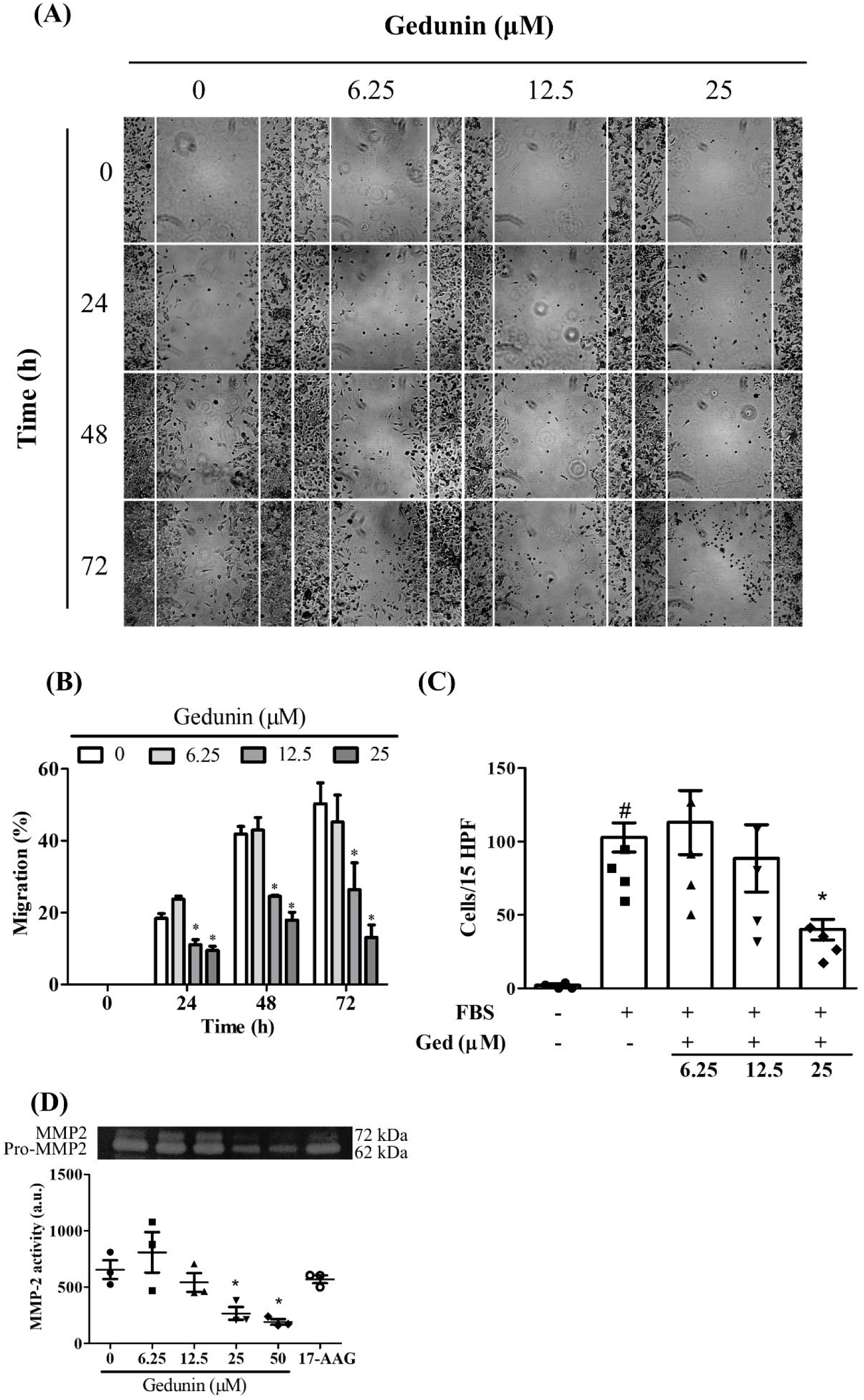
Gedunin impairs in vitro GL261 cell migration and invasiveness. (A) Microscopic views of *the in vitro* scratch assay. GL261 migration into the scratched area 24-72 h after gedunin treatment (6.25-25 μM). (B) Percentages of GL261 cell migration calculated as the ratio of uncovered area (24-72 h) by the initial scratched area (0 h). Results are expressed as the mean ± SEM from two independent experiments (6 well plates). (C) Boyden chamber chemotaxis assay demonstrating the number of gedunin-treated or untreated cells per 15 high power fields (HPF) after 24 h migration. Results are expressed as the mean ± SEM from quadruplicate wells. (D) MMP-2 activity of GL261 cell supernatant assessed by zymography assay 48 h after gedunin treatment (6.25-50 μM). Results are expressed as the mean ± SEM from two independent experiments. * Indicates statistically significant differences (p ≤ 0.05) between treated and untreated cells, whereas # indicates statistically significant differences (p ≤ 0.05) between stimulated and unstimulated cells. FBS: fetal bovine serum.

Extracellular matrix remodeling by the proteolytic activity of tumor cell matrix metalloproteinases (MMPs) is a crucial step for tumor cell invasion in the surrounding tissue [60]. Gedunin also reduced the activity of MMP-2 (Fig. 3D), suggesting that this limonoid impairs different mechanisms associated with tumor cell invasion.

### 3.4 Gedunin modulates signaling proteins in GL261 cells

Ras/ERK is one of the main pro-survival pathways in glioblastoma cells [61,62]. Phosphorylation of ERK 1/2 (pERK) was induced by gedunin (Fig. 4A**-C**) whereas the immunostaining of total ERK1/2 was reduced (Fig. 4D). It is noteworthy that the staining of GAPDH, used as loading control, was also reduced (Fig. 4A; replicates in **Suppl.** Fig. 3) which may be explained by the induction of apoptosis and cell death. GL261 treatment with 17-AAG also reduced total protein content but did not induce ERK phosphorylation within the 24 h incubation.

**Figure 4.**
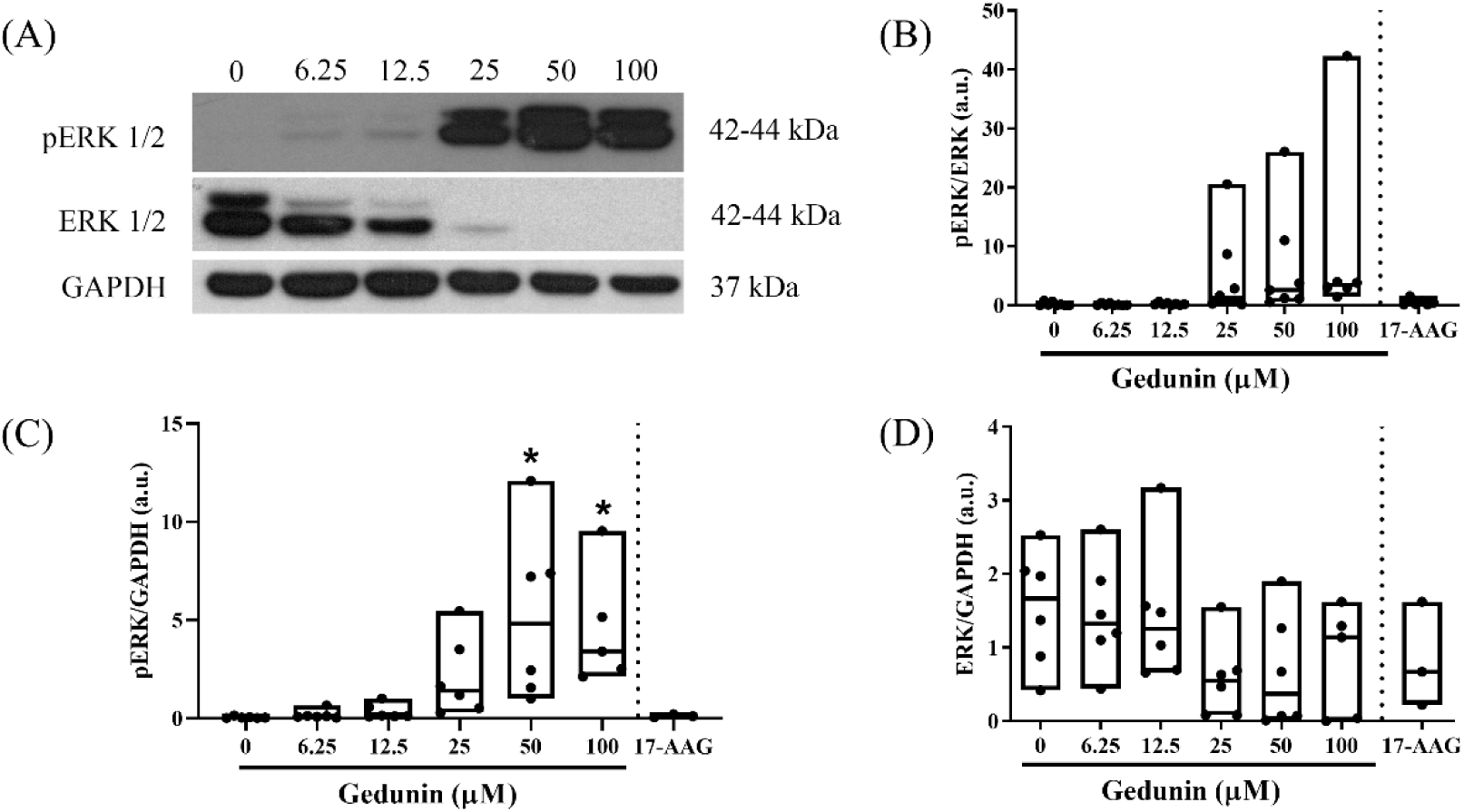
Gedunin activates ERK 1/2 pathway. Western blot analysis of the effect of gedunin (6.25 – 100 μM) on intracellular signaling proteins, 24 h after treatment. (A) Representative immunoblots of one experiment out of 8 independent experiments with a similar trend. Box plots of (B) pERK/ERK, (C) pERK/GAPDH and (D) ERK/GAPDH on GL261 cells. 17-AAG was used as the reference control of HSP90 activity inhibition. GAPDH was used as the loading control. Results are expressed as the mean ± SEM of protein expression from six independent experiments. * indicates statistically significant differences (p ≤ 0.05) between treated and untreated cells. Please find the uncropped blots in supplemental figure 3.

### 3.5 Gedunin effect on glioblastoma in vivo

The orthotopic C57BL/6 mouse glioblastoma model used in the present study has been established as a standard procedure to reproduce human molecular and histological features of glioblastoma [63,64]. Biodistribution data obtained by our group show that ^99m^Tc-labeled-gedunin did not reach substantial levels in the brain tissue when intravenously injected in mice (Costa et al., *manuscript in preparation*). We, therefore, assessed whether gedunin *in vivo* treatment was effective by using osmotic pumps placed *in situ*.

Computed tomography (CT) scans revealed solid tumor growth that occupied an extensive area of the brain tissue of mice from all groups (white mass, top panels in black background of Fig. 5A, indicated by an asterisk in the upper left image scan). Three-dimensional reconstructions of CT images are shown in Figure 5 bottom panels, in which tumor mass is represented by white over gray (brain) areas. Even though tumor volume median values (mm^3^) of gedunin-treated mice were smaller than those for untreated or DMSO-injected mice, variability within groups was high (lowest and highest tumor volume values for each group were: untreated, 21.7 - 75.6; DMSO, 16.8 - 83.6; GED, 11.7 - 67.4). No significant differences were observed between groups (ANOVA, p = 0.106; Fig. 5B, C). **Supplemental** Figure 4 shows CT of mouse brains with and without tumors, for a better illustration of the tumor mass. Body mass (g) was measured before GL261 injection and 28 days later, and no alterations were observed (data not shown).

**Figure 5.**
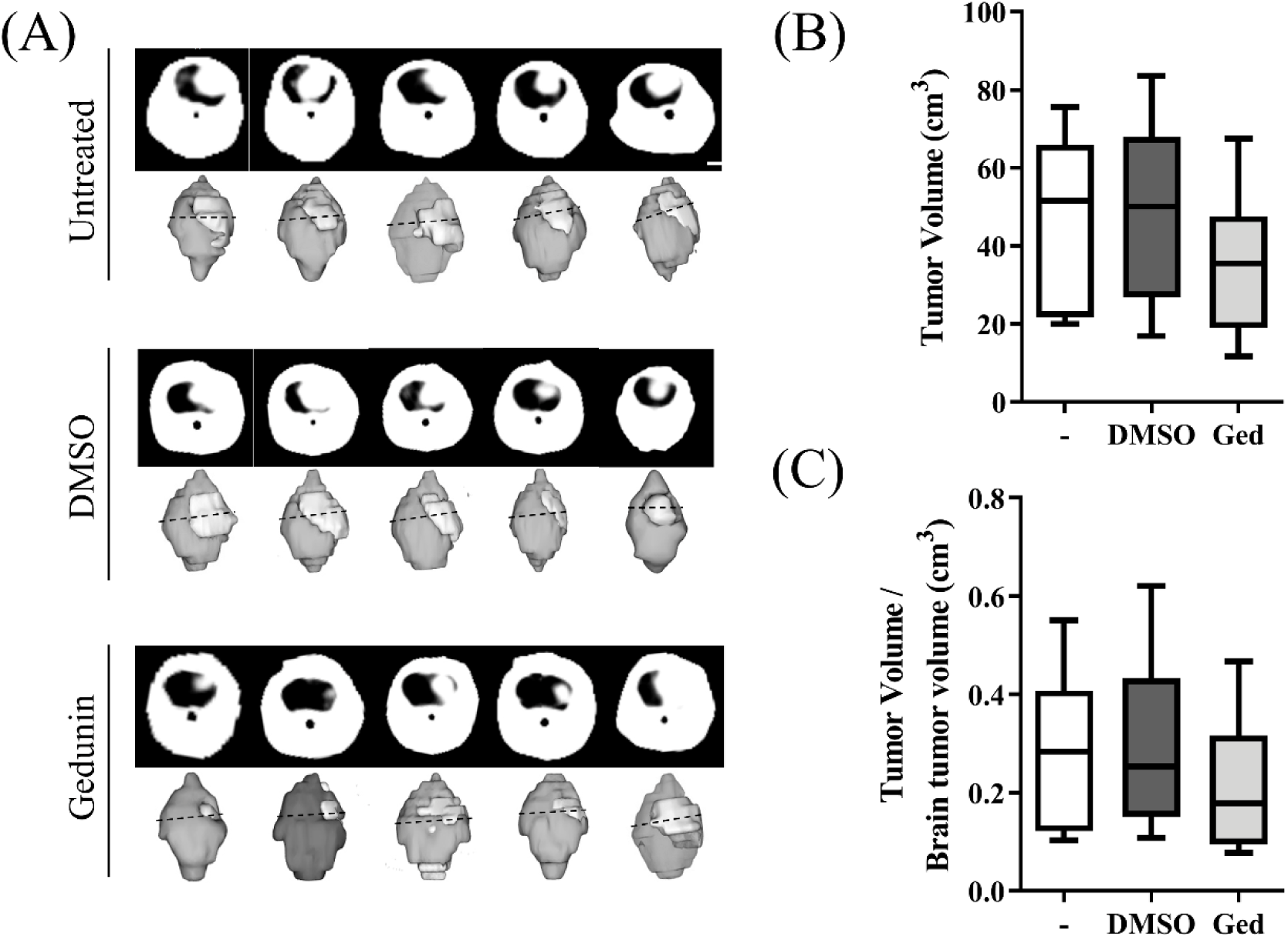
Tumor size analysis of gedunin-treated mice. (A) Representative sequential coronal CT-images and respective 3D brain reconstructions (dorsal view) from untreated, DMSO-injected and gedunin (250 μM)-treated mice. Lighter regions in 3D reconstructions represent brain tumors. Images were acquired after 14 days of *in situ* treatment (Alzet^®^ osmotic minipumps) in GL261 cell-injected mice (1× 10^5^ cells/brain/2 µL). Boxplot graphs show tumor volume values (B) and the ratio between tumor and mouse brain volume values (C) from 6-8 mice per group out of 2 independent experiments. Tumor volumetry was determined by selecting contrast-enhanced regions of interest in coronal sections using InVesalius 3.1.1 software. CT: Computed Tomography. The white area corresponds to the glioblastoma observed in all CT scans. Dashed line: coronal slice cut point. Results are expressed as the mean ± SEM of tumor volume measurements from two independent experiments.

Because clinical CT scanners are designed for imaging human brains, which have volumes roughly 3,000 times larger than those of mice [65–67], the limited spatial resolution may not be precise enough to reliably assess treatment responses in small murine brains [68,69]. To address this limitation, we investigated whether differences between groups could be detected by analyzing tumor area from histological sections. Indeed, Figure 6 illustrates that gedunin-treated mice had reduced tumor central area (see Methods section). In addition, gedunin-treated brain tumors displayed fewer vascular structures (**Table 1** and Supplemental Figure 5).

**Figure 6.**
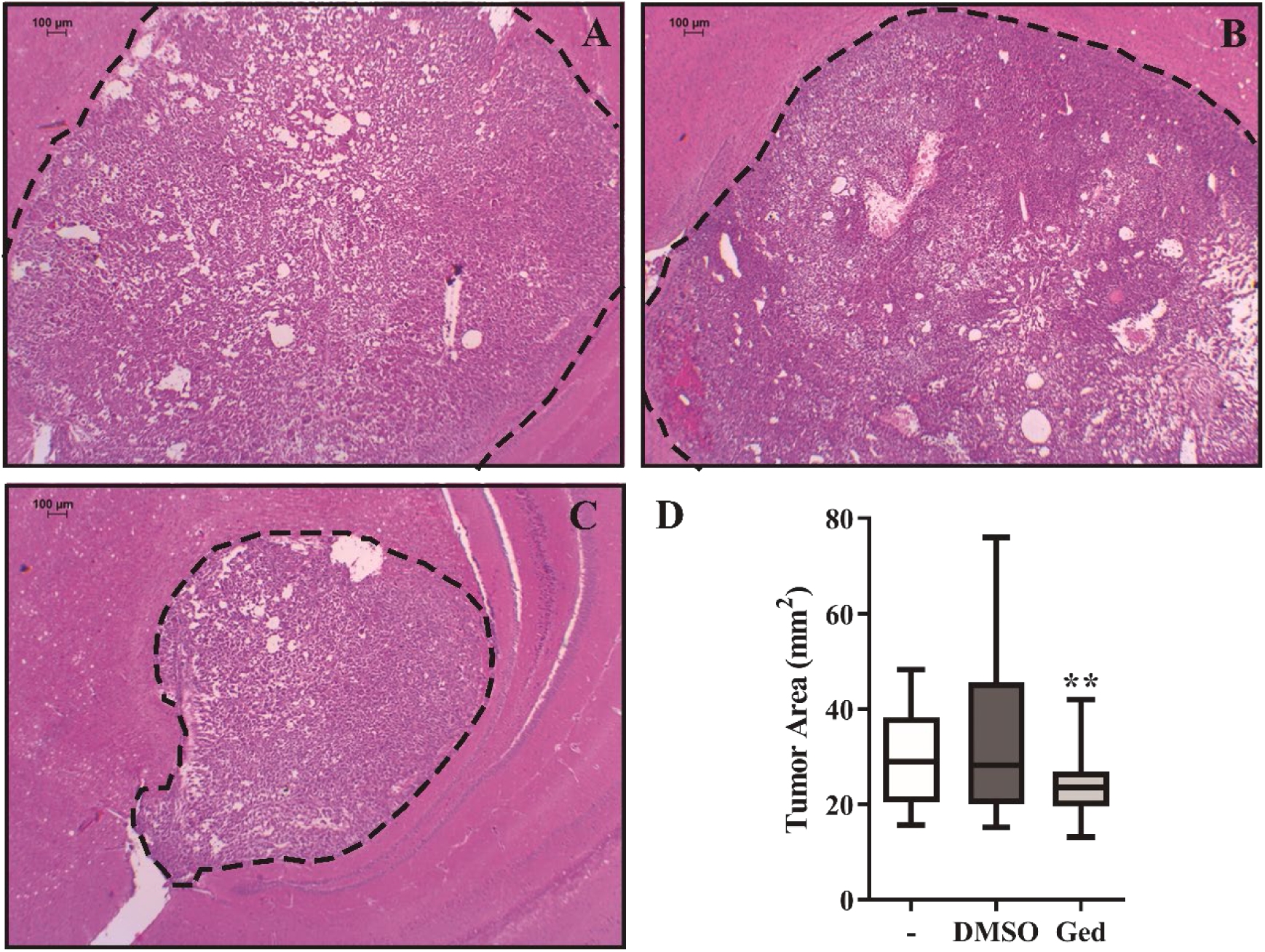
Tumor size analysis of gedunin-treated mice. Representative images of coronal mouse brain slices stained with hematoxylin and eosin showing the tumor central area (100 × magnification) of (A) untreated, (B) DMSO-, and (C) gedunin-treated mice (n=8 per group) 28 days after GL261 cell implantation. (D) Boxplot graph displaying tumor central area values measured in histological brain sections. Results are expressed as the mean ± SEM of tumor central area measurements from two independent experiments. ** Indicates statistically significant differences (p ≤ 0.05) between gedunin-treated and DMSO-treated cells.

**Table 1.**
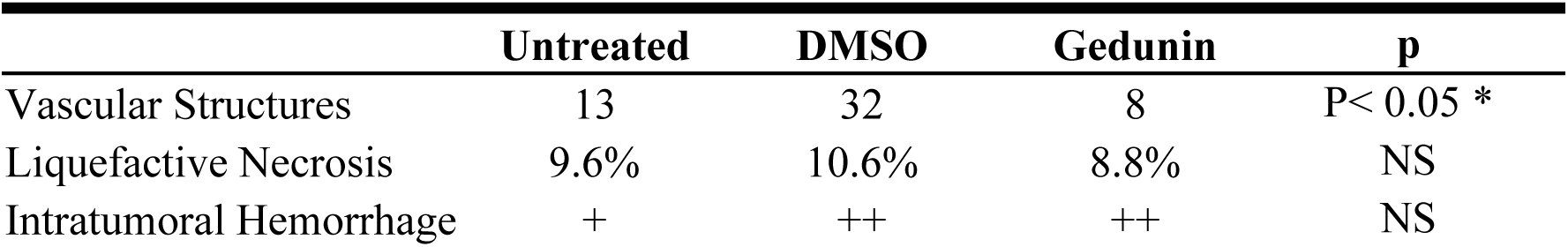
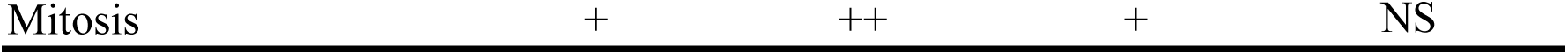
Glioblastoma histology features after gedunin *in vivo* treatment, Table shows absolute and semi-quantitative description of the histological analyses of brain slices and p-values of gedunin-treated vs DMSO-treated mice, analyzed by Tukey’s post-test. NS= non-significant; += discrete; ++= moderate.

|  | Untreated | DMSO | Gedunin | p |
| --- | --- | --- | --- | --- |
| Vascular Structures | 13 | 32 | 8 | $P < 0.05$ * |
| Liquefactive Necrosis | 9.6% | 10.6% | 8.8% | NS |
| Intratumoral Hemorrhage | + | ++ | ++ | NS |
| Mitosis | + | ++ | + | NS |

## 4. Discussion

Uncontrolled proliferation and apoptosis resistance are central features of tumor development, being major targets for cancer treatment [19]. HSP90 is a key factor for tumor growth, since it chaperones several essential proteins involved in oncogenic processes. In the present study, we demonstrate that gedunin, a natural occurring HSP90 inhibitor, impairs murine glioblastoma cell growth *in vitro*, by arresting proliferation and inducing cell death in non-toxic concentrations to primary mouse brain cells. As previously shown in human cell lines [43,44], gedunin reduced murine glioblastoma cell migration, tumor-related signaling pathways, and extracellular metalloproteinase MMP-2 activity, features involved in glioblastoma growth. GL261-orthotopically injected C57BL/6 mice treated with gedunin had reduced presence of vascular structures, an important hallmark of tumor maintenance and growth.

Gedunin and other HSP90-binding agents induce the expression of HSP70 in cancer cells due to HSP90 inhibition (**Suppl.** Figure 6) [70–72] and show antiproliferative and cytotoxic effects against different tumor cells [37,39–42]. HSP90 inhibitors have increased affinity to the HSP90 machinery expressed by tumor cells [16]. This supports our data showing that gedunin toxicity against GL261 cells was achieved at a concentration lower than half of the required against primary glial cells [73]. Gedunin-triggered apoptosis, detected by phosphatidylserine exposure on the cell surface, was confirmed by flow cytometric analysis of cell physical criteria that showed the reduced size and increased granularity of apoptotic cells, due to the formation of apoptotic bodies and plasma membrane blebbing [55]. Our results clearly show that, as apoptosis progresses, GL261 cell FSC becomes progressively lower (likely due to cell dehydration and shrinkage), whereas SSC increases, due to cytoplasm and nuclear condensation [55,57].

Glioblastoma is a highly invasive and diffuse tumor that requires cell migration and extracellular matrix degradation to spread to adjacent tissue [74,75]. Scratch and chemotaxis assays showed that gedunin diminished GL261 migration, in concentrations and time points lower than the ones that induced cytotoxicity. Another way by which gedunin controls tumor invasiveness is by inhibiting MMP-2 activity, with a higher effect than 17-AAG. Indeed, HSP90 stabilizes and promotes the activation of extracellular MMP-2, being crucial for GL261 and HT-1080 tumor cell invasion [76,77].

Although total ERK ½ was reduced, gedunin promoted a strong sustained (24 h) ERK1/2 activation/phosphorylation, corroborating previous work which demonstrate that HSP90 inhibition can also prompt the phosphorylation/activation of ERK [78,79]. The activation of ERK ½ pathway is associated with mechanisms involved in GBM development such as proliferation, invasion, metabolism, and death [80,81]. However, depending on the cell type and stimulus, ERK ½ sustained activation can promote cell cycle arrest, autophagic vacuolization, or caspase activation leading to cell senescence, autophagy, or apoptosis, respectively [82–84], mechanisms that might be involved in the reduced levels of GL261 cell proliferation, migration, and induced programmed cell death observed in this work.

As previously mentioned in the results section, even though our findings indicate that gedunin fits the predicted ADMET criteria for druggability and BBB permeation, systemic (i.v.) administration of ^99m^TC-labeled gedunin failed to reach the brain tissue at substantial levels (Costa *et al*., manuscript in preparation). Hence, we used gedunin-filled Alzet^®^ minipumps to perform *in situ* treatment (resembling the FDA-approved Gliadel^®^ [carmustine] implants) to ensure that gedunin would be available at the tumor site. It is important to note that the residual volume of gedunin inside the minipumps was measured at the end of treatment to ensure that gedunin was released *in situ* (values shown in **Suppl. Table 1**). It is important to emphasize the limitations of the CT mouse brain image acquisition using a clinical CT scanner, since the limited spatial resolution of clinical CT scanners might not be precise enough to analyze such a small brain [68,69]. While our findings reinforce the feasibility of using clinical CT scanners for *in vivo* brain tumor imaging in mice, their use for detecting treatment responses in tumor volume requires further investigation. Another limitation of our study is that mouse tumor sizes were not evaluated over time, since CT scan was only feasible in mice with no minipump implants, due to the stainless steel of brain infusion cannulas.

As expected, GL261-injections triggered the development of mouse glioblastoma in orthotopically-injected C57BL/6 mice with liquefactive necrosis, hemorrhage, mitosis, and increased vascular structures [63]. *In situ* treatment with gedunin significantly reduced tumor central area and the presence of vascular structures, which might decrease tumor nutritional supply.

The severity and the poor prognosis of patients bearing glioblastoma call for new therapeutic targets and drugs. Data presented in this paper sheds light on alternative treatments for this type of tumor by revealing GL261 dependency of HSP90 and gedunin as a potential lead compound.

## Supporting information

Costa et al. - Supplemental Figures and Table - BioRxiv - 11-09-25 - revised

## List of abbreviations

17-AAG: 17-(Allylamino)-17-demethoxygeldanamycin/tanespimycin
AKT/PKB: protein kinase B
ARRIVE: Animal Research: Reporting of *In Vivo* Experiment
BCR/ABL: breakpoint cluster region/Abelson murine leukemia virus
CFSE: carboxyfluorescein succinimidyl ester
DMEM: Dulbeccós modified Eaglés medium
DMSO: dimethyl sulfoxide
EDTA: ethylenediamine tetraacetic acid
Erk ½: extracellular signal-regulated kinases ½
FBS: Fetal bovine serum
FITC: fluorescein isothiocyanate
GAPDH: glyceraldehyde-3-phosphate dehydrogenase mouse
HER2/neu, ErbB2: human epidermal growth factor receptor 2
HPF: high-power fields
HSP90: Heat Shock Protein 90
IgG: immunoglobulin G
MMP: matrix metalloproteinase
MTT: 3-(4,5-dimethyl-2-thiazolyl)-2,5-diphenyl-2H-tetrazolium bromide p44/42
MAPK: mitogen-activated protein kinase
PBS: phosphate buffer saline
PI: propidium iodide
SDS-PAGE: sodium dodecyl sulfate-polyacrylamide gel electrophoresis
Src: proto-oncogene c-Src

## Author contributions

CP, TEMMC, and TEK conceived and designed the work. TEMMC, CP, TEK, ACM, and MGH designed experiments. TEMMC, CP, TAP, LNS, EMC, CCF, CMMM, JXP, VGR, YAV, and TEK performed the experiments, acquired, and/or analyzed data. CP, TEK, ACM, JXC, and MGH contributed with funding and resources. CP supervised the work. CP and TEMMC wrote the original draft. CMMM and TEK wrote and revised the manuscript. All authors revised and edited the final manuscript.

## Acknowledgments

The authors gratefully acknowledge Dr. André Luis Peixoto Candéa, D.Sc. Erika Tatiana Muñoz, Dr. Samara Cristina Ferreira Machado, Edson Santos Silva, Filipe Leal Portilho, Ubirajara Ribeiro, and Maria Heloisa Oliveira for technical support. We also thank the Institute for Biomodels Science and Technology (Fiocruz) for providing mice, and the Center for Cancer Control (CUCC, UERJ) for the help with CT imaging acquisition.

