## Supplementary material for "Effect of gedunin on glioblastoma proliferation and invasiveness: *in vitro* and *in vivo* approaches": Costa et al. - Supplemental Figures and Table - BioRxiv - 11-09-25 - revised

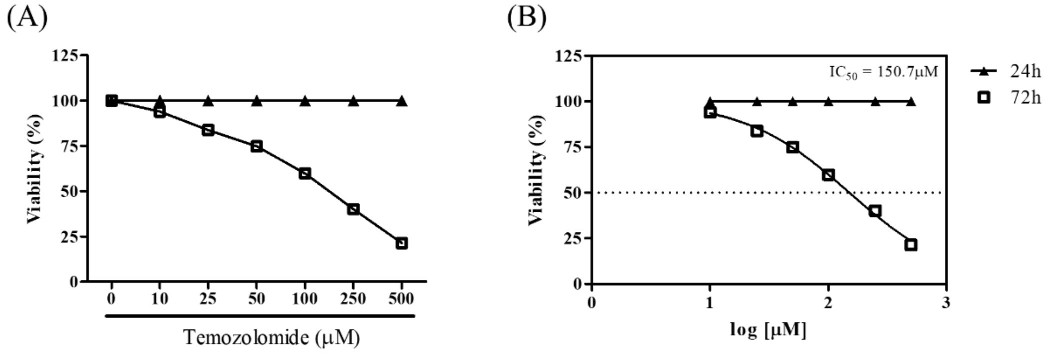


**Supplemental Fig. 1 -** *Temozolomide induces time- and concentration-dependent GL261 death.* GL261 cells were incubated with temozolomide (10 – 500 μM in 5 % DMSO) for 24 h and 72 h (37 °C, 5 % $CO_2$). Viability was evaluated by MTT reduction assay (SpectraMax M5 – Molecular Devices) and results are expressed as the mean ± SEM of percentage of viability (A) and inhibitory concentration (B) from quadruplicate wells from one experiment (96 well plates, flat bottom).

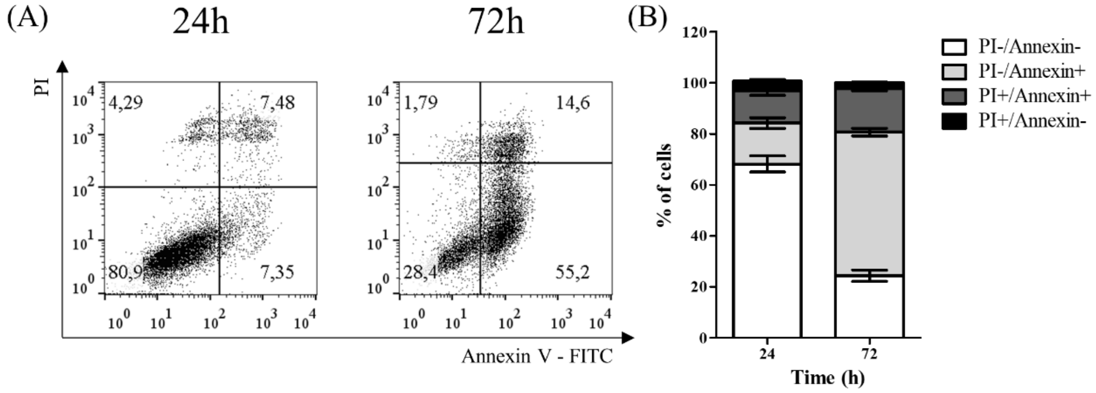


**Supplemental Fig. 2 -** *Effect of 17-AAG on GL261 cell apoptosis*. GL261 cells were incubated with 17-AAG (1 μM) for 24h and 72h (37 °C, 5 % $CO_2$) and labeled with annexin V-FITC and PI for flow cytometric analysis (FACScalibur, BD). (A) Representative dot plots of FlowJo analysis and (B) bar graph expressing mean ± SEM of triplicate wells from three independent experiments (96 well plates, flat bottom).

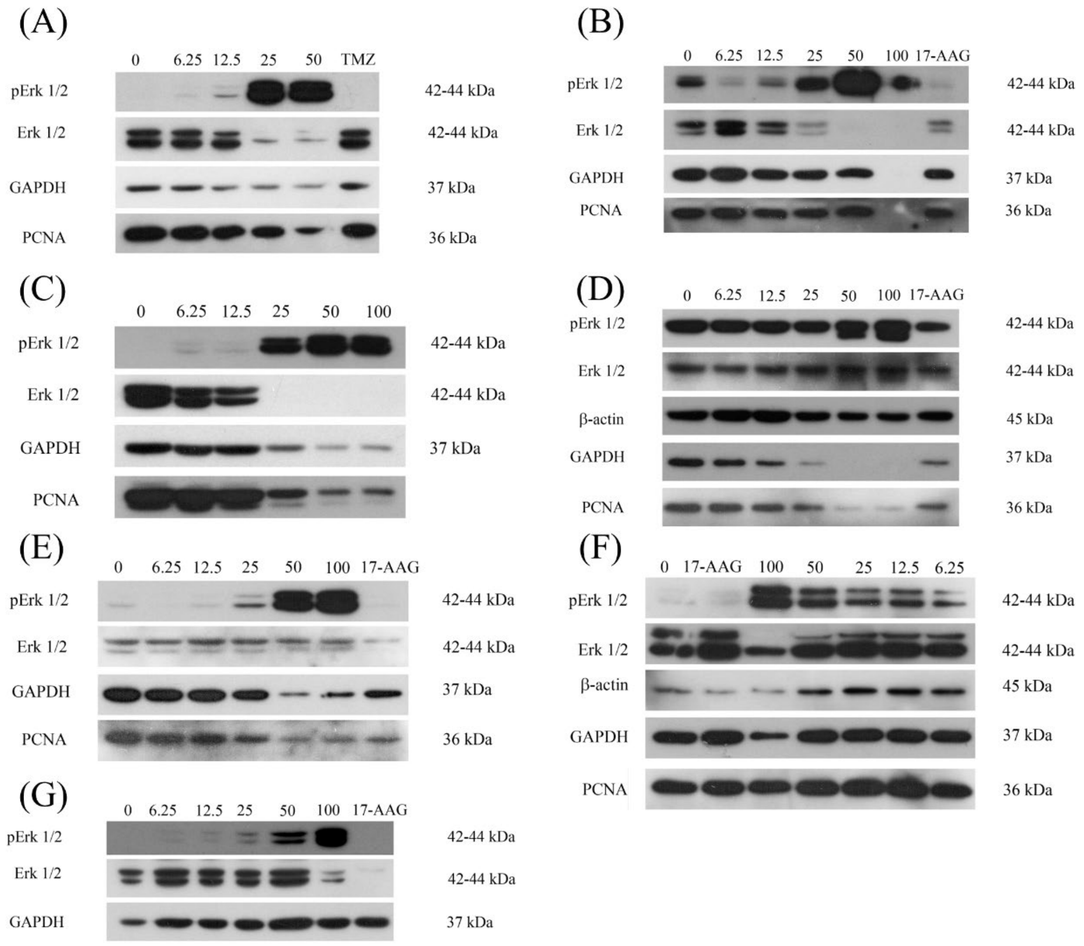


**Supplemental Fig. 3 –** *Western blots of the effect of gedunin on intracellular signaling proteins 24 h after treatment.* Panels show immunoblots from seven independent experiments (A-G). 17-AAG (1 μM) was used as the control of HSP90 activity inhibition. GAPDH was used as the loading control. Please find the uncropped blots at the end of this supplemental file.

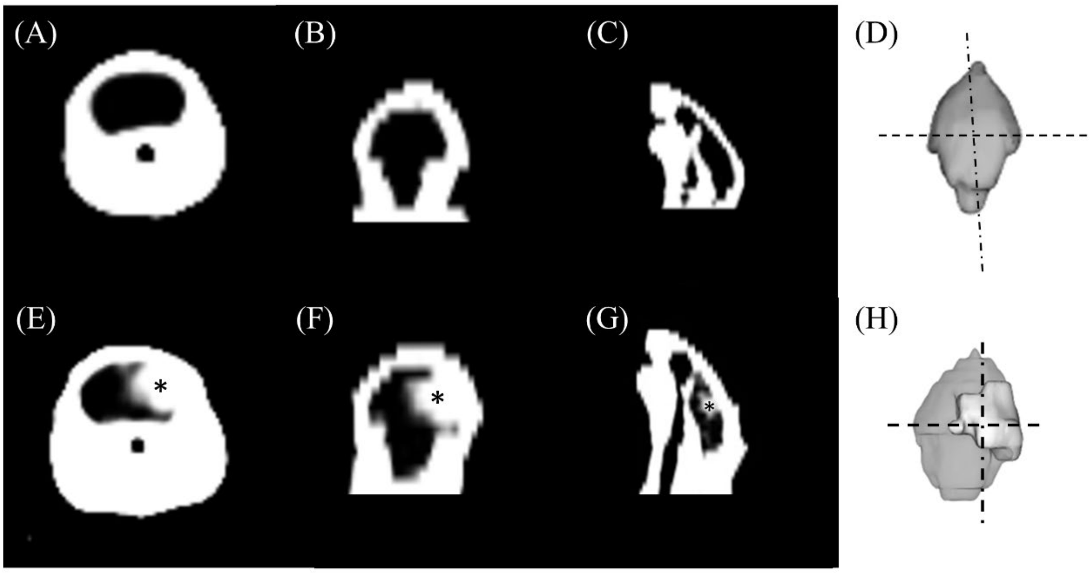


**Supplemental Fig. 4 -** *Brain CT images of C57BL/6 mice.* (A-C, E-F) Representative CT-scans at different orientation planes: coronal (A, E), axial (B, F), and sagittal (C, G) of two C57BL/6 mice, one without (A-C) and another with tumor (E-G, *glioblastoma). (D, H) Top view of 3D brain reconstructions from the same animals depicted in (A-C, E-F). Lighter region in H shows the extent of the brain tumor. Horizontal dashed line: coronal slice cut point; vertical dot-dashed line: sagittal slice cut point.

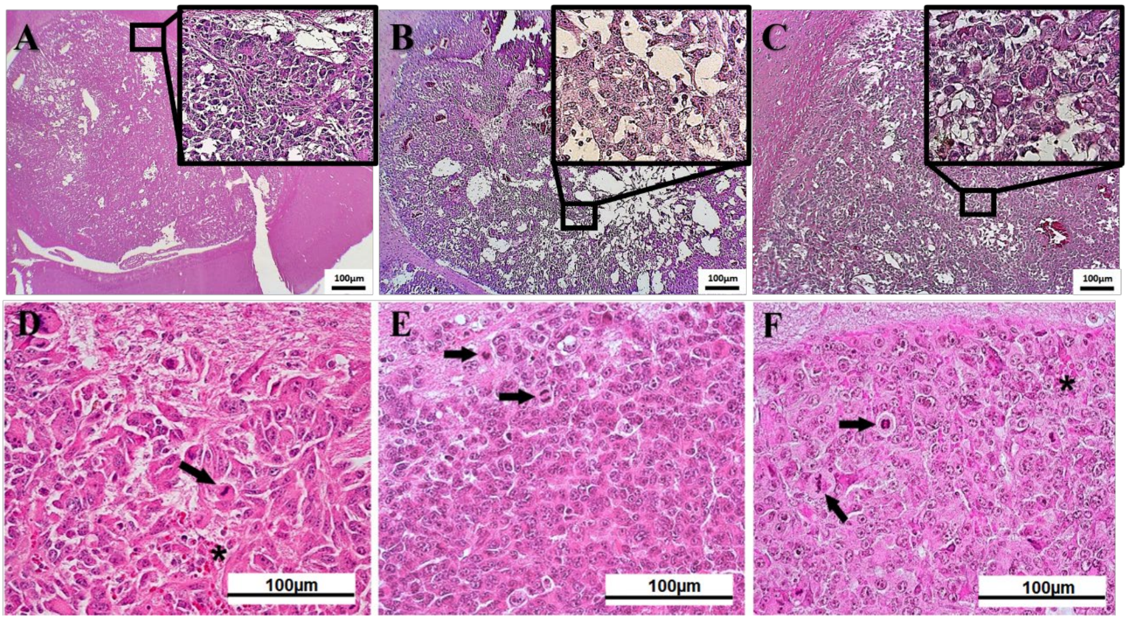


**Supplemental Fig. 5.** *Histopathological analysis of gedunin-treated mice.* Representative images of coronal brain slices from stained with hematoxylin and eosin 28 days after GL261 cell implantation of untreated (A, D), DMSO- (B, E), and gedunin-treated (C, F) mice (n=8 per group). Upper panels (A-C) depict low magnification images (40 ×) of the extent of the tumor area for each experimental group. Corresponding high-magnification inserts (400 ×) of intratumoral perinecrotic regions show liquefactive necrosis (white regions). Lower panels (D-F), high magnification images (1000 ×) from the same animals in (A-C), show mitosis (black arrows) and areas of hemorrhage (asterisks) in intratumoral regions.

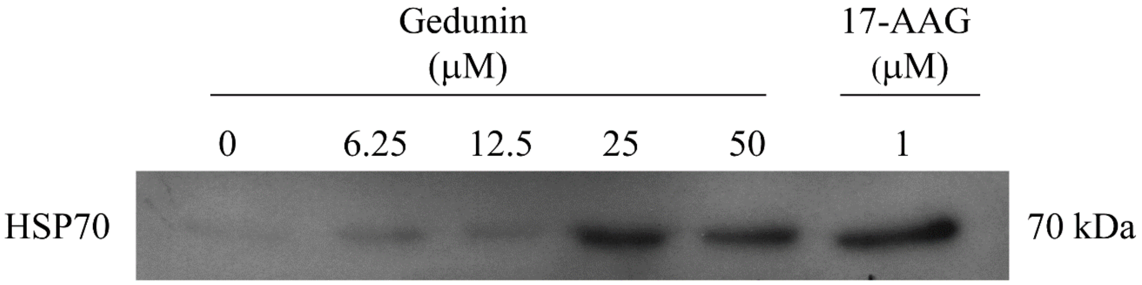


**Supplemental Fig. 6 -** *Gedunin induces HSP70 secretion.* GL261 cell supernatant was collected 48 h after treatment, lyophilized, and resuspended in PBS for western blot analysis. Immunoblot shows the HSP70 staining from one experiment with one replicate. Protein loading (20 μg/lane) for SDS-PAGE was determined after Lowry protein method.

**Supplemental Table 1. Osmotic minipump content release**

| Treatment | Volume released by the osmotic minipumps per day (μL/day) |
|---|---|
| DMSO | 4.2 ± 0.2 |
| Gedunin | 3.9 ± 0.2 |

Alzet® osmotic minipumps were filled with DMSO (2.5 %) or gedunin (250 μM; 1.2 μg/μL). The volume released per day was calculated by subtracting the final volume (obtained on the day of minipump removal) from the initial volume in each minipump, considering the number of days between minipump filling and the end of the treatment. Data are represented as the mean ± SEM of each treatment (n = 8/group).

**Western blot original uncropped images**
**Figure 4A**

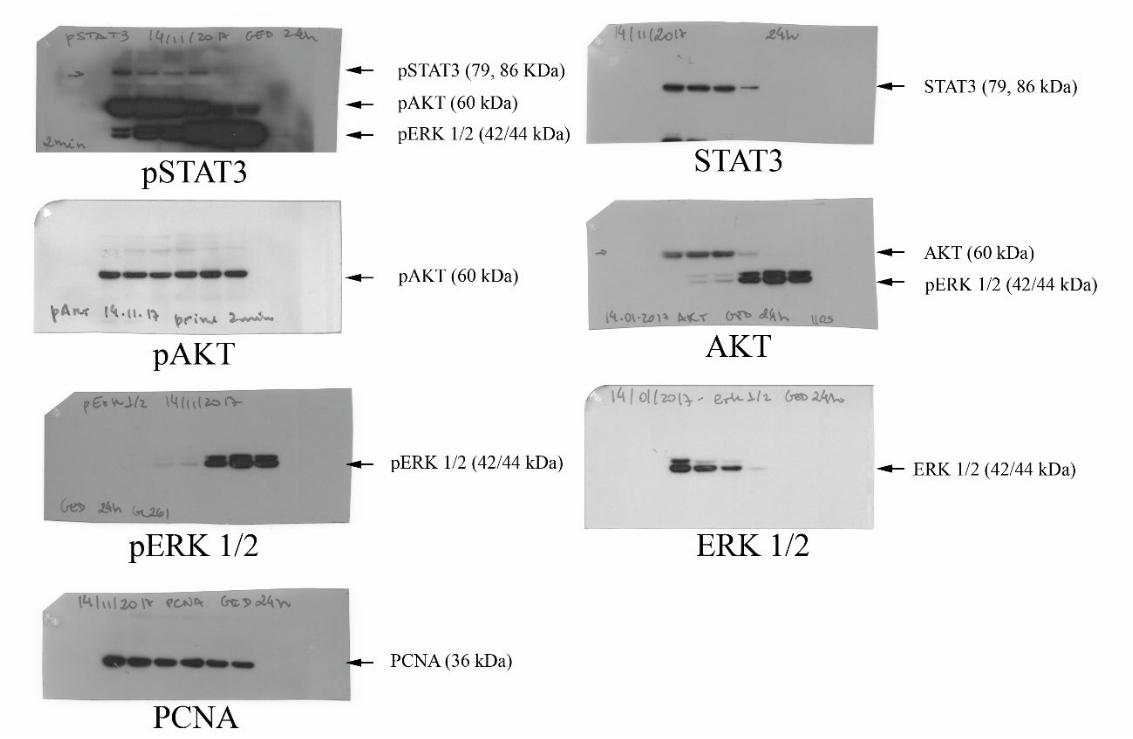


**Supplemental Figure 3A**

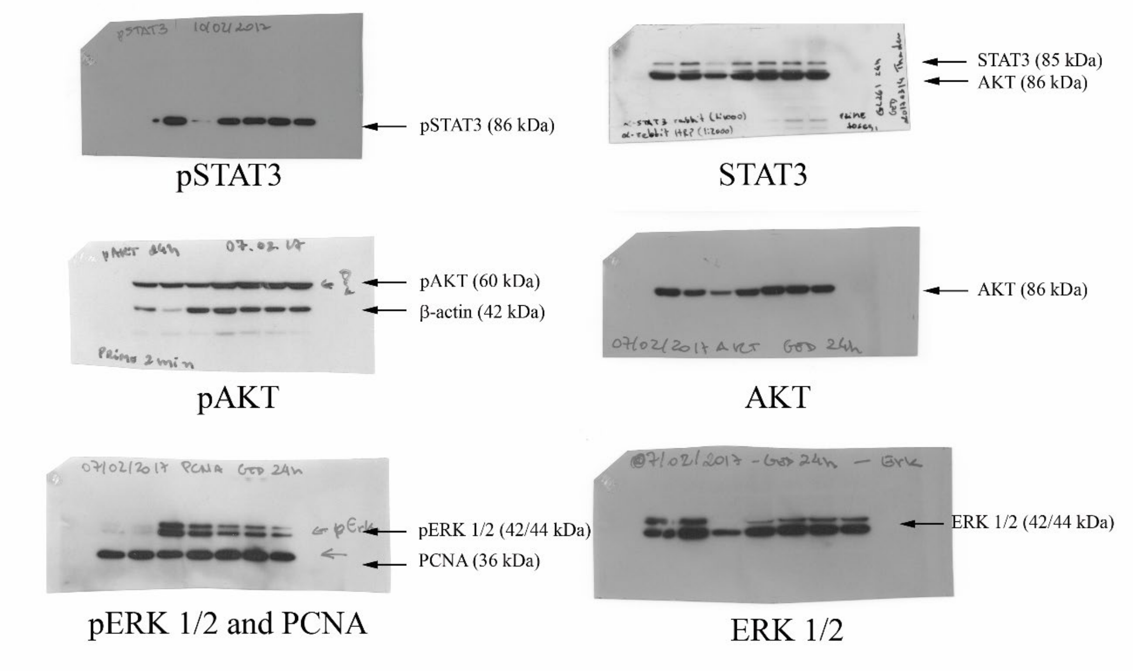


**Supplemental Figure 3B**

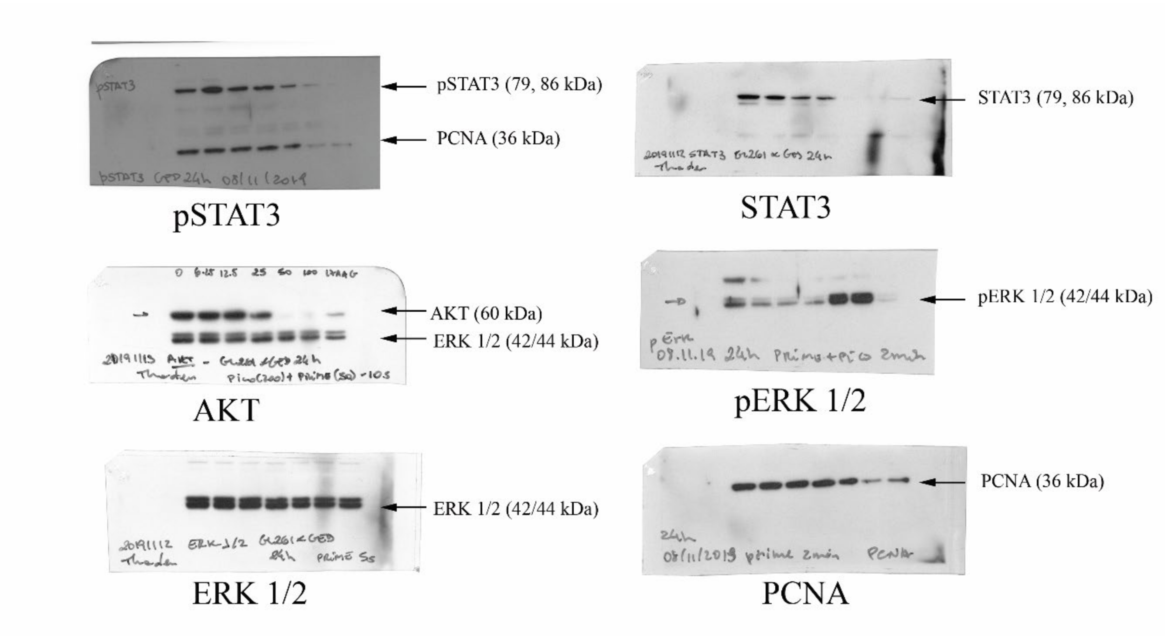


**Supplemental Figure 3C**

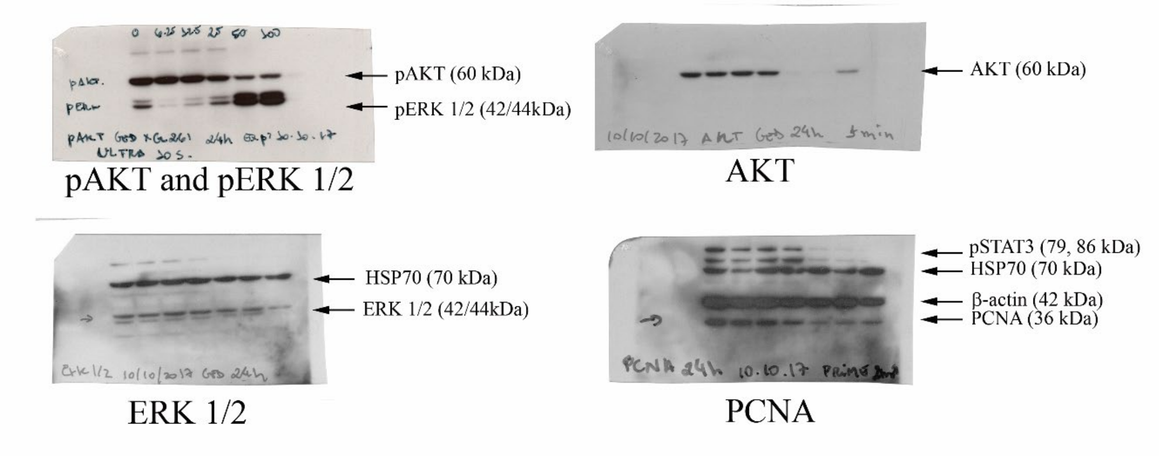


**Supplemental Figure 3D**

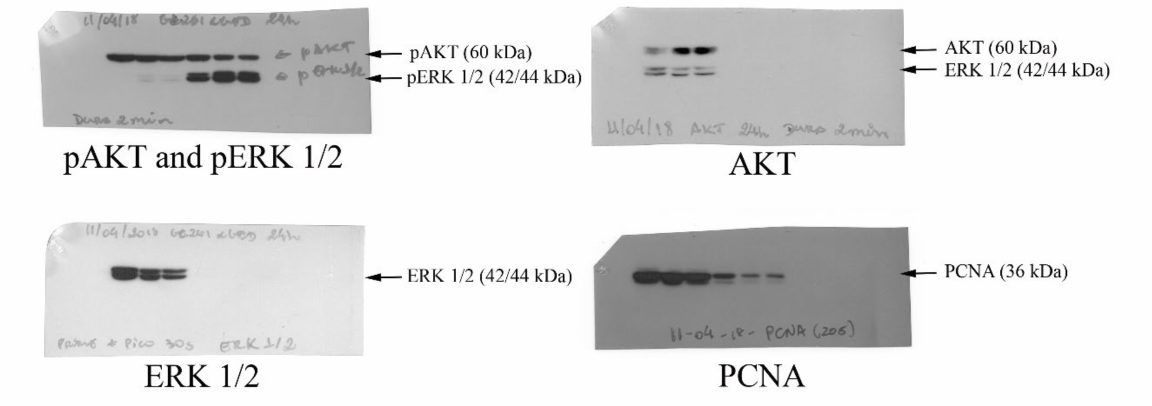


**Supplemental Figure 3E**

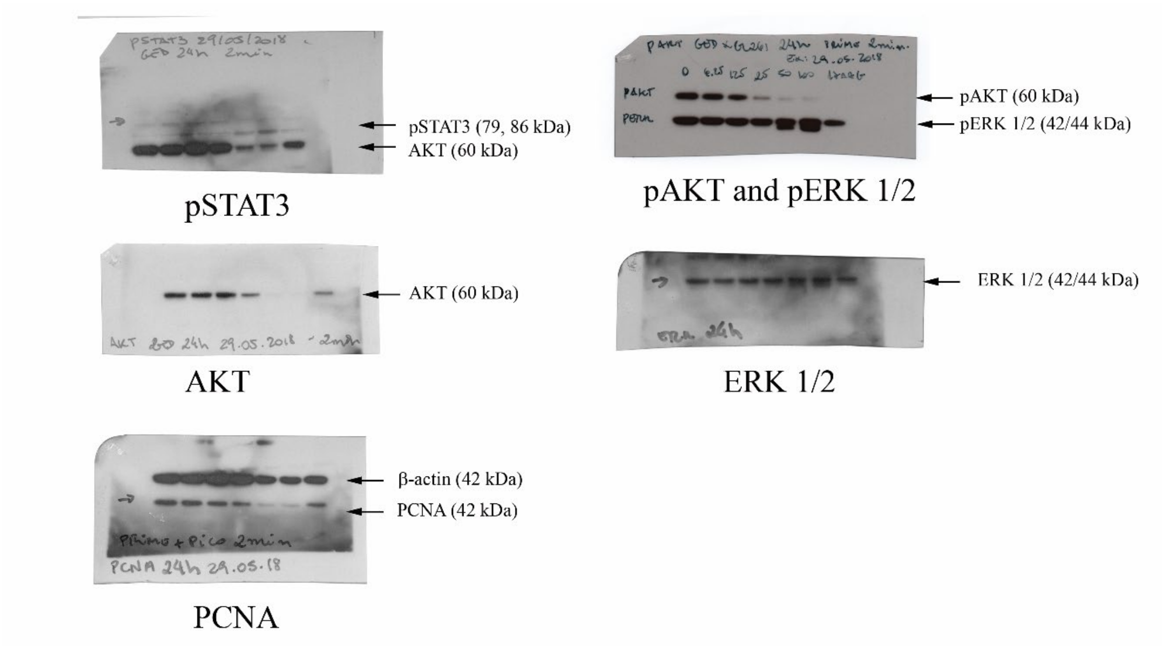


**Supplemental Figure 3F**

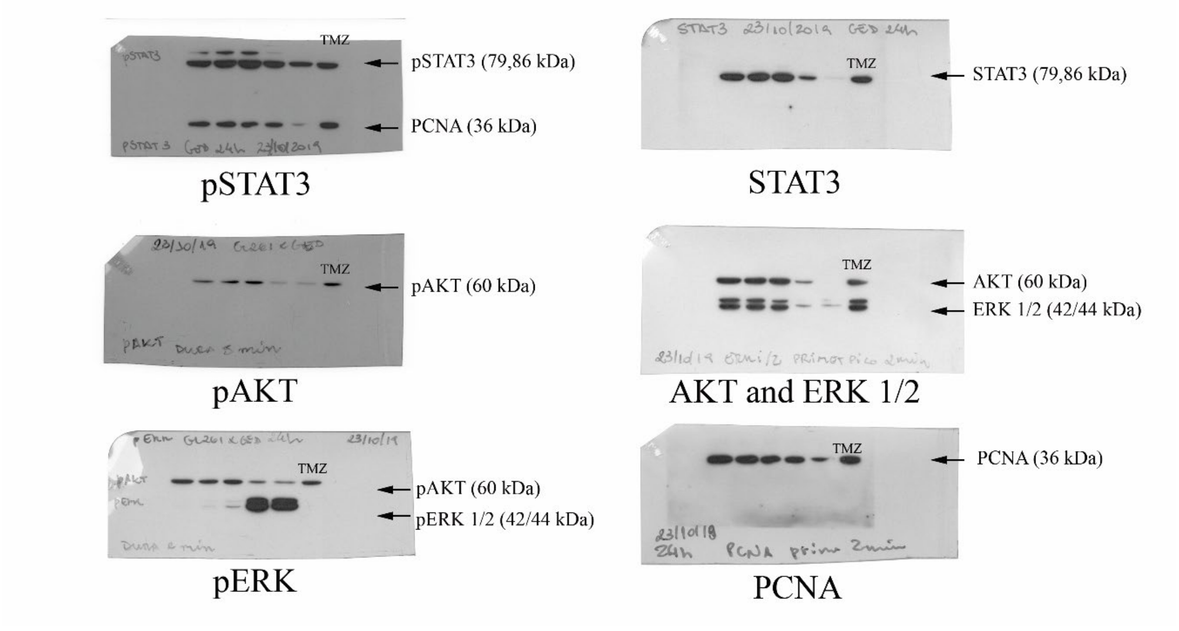

**Supplemental Figure 3G**

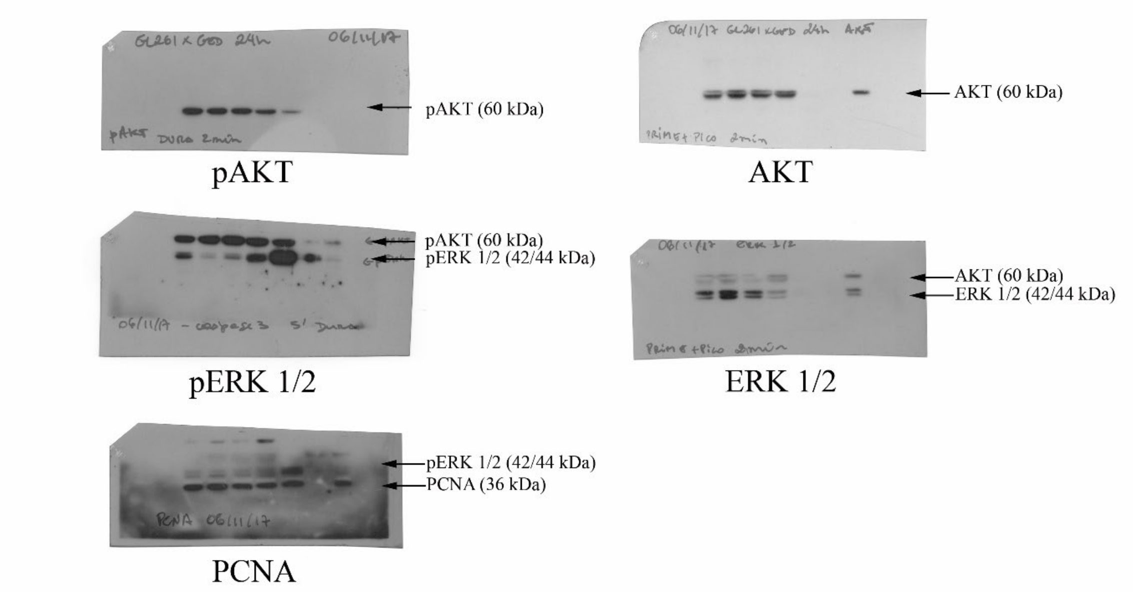
